# A novel plasmid-transcribed regulatory sRNA, QfsR, controls chromosomal polycistronic mRNAs in *Agrobacterium tumefaciens*

**DOI:** 10.1101/533729

**Authors:** Benjamin Diel, Magali Dequivre, Florence Wisniewski-Dyé, Ludovic Vial, Florence Hommais

**Author notes:** Corresponding author: Florence Hommais, Microbiologie Adaptation Pathogénie, Université Lyon 1, F-69622 Villeurbanne cedex, France.

## Abstract

Plasmids are mobile DNAs that adjust host cell functions for their own amplification and dissemination. We identified QfsR, a small RNA, transcribed from the Ti plasmid in the phytopathogen *Agrobacterium fabrum*. QfsR is widely conserved throughout RepABC plasmids carried by *Rhizobiaceae*. Target prediction, expression analysis and site-direct mutagenesis experiments show that QfsR directly pairs within polycistronic mRNAs transcribed from chromosomes (involved in flagella apparatus and succinoglycan biosynthesis) and Ti plasmid (involved in conjugative transfer). QfsR leads to a coordinated expression of whole polycistronic mRNA molecules. Whereas a lack of QfsR induces motility and reduces succinoglycan production, its overproduction increases the quorum sensing signal accumulation and the Ti plasmid conjugative transfer. Based on these observations, we propose QfsR as a hub connecting regulatory networks of motility, succinoglycan biosynthesis and plasmid conjugative transfer. To our knowledge, QfsR is the first example of a plasmid-encoded sRNA that controls chromosomal polycistronic mRNAs.

**Significance:** Plasmids represent an important cost for the hosting cell although some are beneficial under certain circumstances. *Agrobacterium tumefaciens* harboring Tumor inducing plasmid (pTi) are able to infect plants and to use specific resources produced by the infected cells. We characterized QfsR, a novel small RNA (sRNA) from pTi, that directly regulates plasmid polycistronic mRNA but also chromosomal ones. QfsR contributes to a fine-tuned regulation of bacterial motility, exopolysaccharide biosynthesis and conjugative dissemination of pTi. Our results report the first plasmid-encoded sRNA able to modify and coordinate cellular behaviour probably for the benefit of the plasmid dissemination and tight crosstalk between plasmid and chromosome. This could be widespread since QfsR homologs were predicted in other plasmids of *Rhizobiaceae* symbionts and pathogens.

## Introduction

Plasmids are foreign DNAs whose expression and replication can impose a significant cost on host cells. However, their acquisition might be beneficial under certain circumstances, conferring advantageous traits, such as antibiotic resistance, ability to catabolize nutrients, and pathogenesis (1)(2)(3). To reduce fitness cost, both plasmid and host cells have developed a tight regulatory network, which may involve crosstalk between chromosome and plasmid. Indeed, several chromosomal regulators control the expression of plasmid functions (4)(5) and more rarely plasmid regulators control chromosomal genes (6)(7)(8).

*Agrobacterium* with the Tumor inducing (Ti) plasmid is responsible for bacterial virulence on a variety of dicotyledonous plants, as it induces the production of plant growth hormones that cause cell proliferation (tumours) (9). The major virulence factors encoded by the Ti plasmid include a type IV secretion system and accessory proteins (TSS4 and Vir) responsible for T-DNA transfer and integration into the plant genome (10). In addition, opines produced by transformed plant cells are catabolised by pTi-harboring agrobacteria, giving rise to an agrobacteria-specific ecological niche (11). Opines also act as signals promoting pTi conjugal transfer (12). Conjugal transfer involves a second Ti plasmid-encoded T4SS, the Tra/Trb complex, whose production is regulated by quorum sensing signals (12)(13). Even though the ability to colonize the plant and the presence of a particular ecological niche are beneficial traits encoded by the Ti plasmid, their expression is costly and only relevant under certain conditions. Hence, there is a complex regulatory network that tightly controls Ti plasmid expression and replication to counterbalance the fitness cost of plasmid carriage (14)(15)(16)(17). Some chromosomal regulators are involved i.e. ChvE and ChvIG (4)(5) but to date, no plasmid regulator has been identified in this crosstalk between plasmid and chromosome.

Small regulatory RNAs have been identified in the past years as posttranscriptional regulators (18). These regulatory RNAs generally control mRNA translation and stability by direct RNA-RNA base pairing. Among them, *trans-acting* RNAs require a short stretch of sequence complementarity to be sufficient for regulation. Base pairing could also require the assistance of the Hfq RNA chaperone (19). Recently, RNA-seq analyses have revealed sRNA repertoires of *Agrobacterium fabrum* C58, formerly called *A. tumefaciens* C58 (20)(21)(22). We previously identified 75 candidates transcribed from the Ti plasmid of C58 strain (pTiC58) and a large majority of them have a constant expression level whatever the conditions tested (22).

Here, we report on the identification and characterization of one candidate, named QfsR. We determined the functions of QfsR through the identification of three of its targets that are large polycistronic mRNAs: the conjugal transfer operon from the Ti plasmid, the flagellar operon from the circular chromosome and the succinoglycan biosynthesis operon localized on the linear chromosome. We showed that QfsR regulates mRNA targets by interacting directly with at least one short base-pairing site per polycistronic mRNA, apparently coordinating expression level of all the polycistrons. QfsR is the first example of a trans-regulatory sRNA transcribed from a plasmid that directly modulates chromosome-encoded mRNAs.

## Results

### QfsR is transcribed from the Ti plasmid

Preliminary RNA-seq experiments assigned RNA1083 as a putative *trans-encoded* RNA transcribed from the minus strand of the pTiC58 between genes *atu6119* and *atu6120*, a region upstream of a conjugative gene cluster *(tra* polycistrons) (22). The determination of its transcriptional start and stop revealed a small transcript of 188 bases in length beginning at base 139,262 and ending at base 139,075 (Fig. S1). Its predicted secondary structure is robust, strong and stable (Fig. S1). We renamed it QfsR (**Q**uorum sensing **f**lagella and succinoglycan biosynthesis **R**NA regulator), with reference to the phenotypes modulated by this sRNA (see below).

**Fig. S1.**
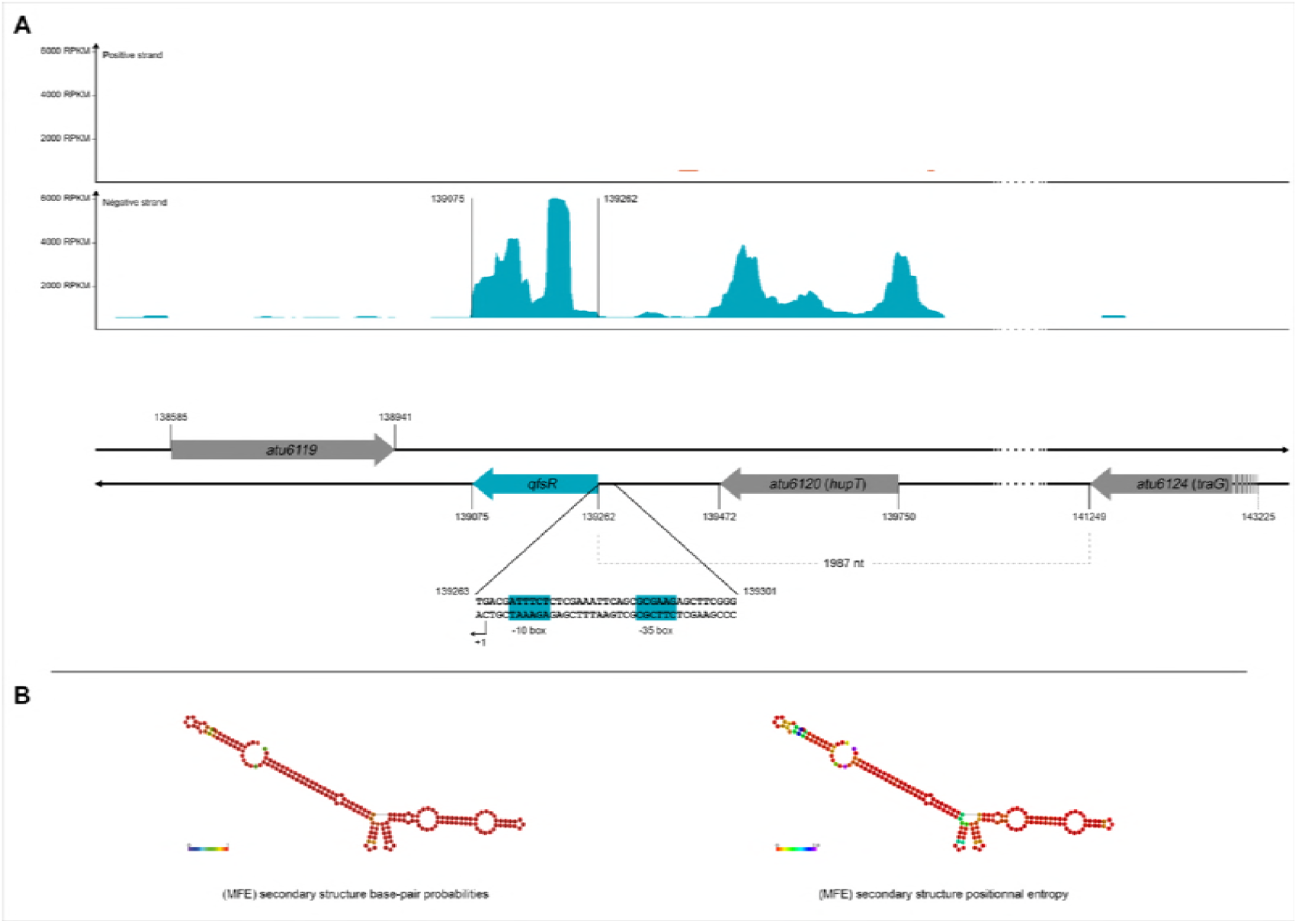
Genetic organization of the *qfsR* locus from the Ti plasmid and predicted secondary structure of the transcribed sRNA. A) Data from *A. fabrum* sRNA-seq displayed using Artemis (22). Strand-specific coverages against the reference strain are displayed as two plots: at the top is the coverage displayed from the plus strand and the bottom plot shows coverage from the minus strand. The genome annotation is displayed underneath. The genomic environment of the *qfsR* locus is shown. Arrows correspond to genes. The 5’-end and 3’-end of the *qfsR* gene were determined from RACE-PCR results. Sequences in blue boxes correspond to the putative promoter region of *qfsR -* the -10 box (AGAAAT, with consensus TATNNT) and the - 35 box (CTTCGC with consensus CTTGNN). +1 represents transcription start site of *qfsR*. (RPKM: Reads Per Kilobase of transcript per Million reads mapped). B) The secondary structure of QfsR was predicted using the RNAfold algorithm (Minimum Free Energy) and results obtained are colored according to base-pair probabilities (on the left) or positional entropy (on the right).

### QfsR is widely conserved among RepABC plasmids carried by Rhizobiaceae

The localization of *qfsR* gene on pTiC58 questioned its conservation. Therefore, we investigated whether QfsR homologs could also be detected within genomes available in the NCBI Nucleotide collection (nt) and in the MicroScope genome database (23). Using the BLASTN algorithm, we identified 395 candidates showing various sequence similarities with *qfsR*. Using RNAfold algorithm, we predicted the secondary structures of these candidates and compared them using the RNAforester algorithm (24). We selected 230 putative QfsR homologs according to the high conservation of their secondary structure. 88 candidates, most of which belonging to *Agrobacterium* genomes (Figure 1A), present also a high percentage of sequence identities and a good sequence coverage and 142 candidates present a high conservation of their secondary structures but a low conservation level of their nucleotide sequences (Table S1, Figure 1B). Structural homologs of QfsR were distributed mostly on plasmids for the following *Rhizobiaceae: Agrobacterium, Rhizobium* and *Sinorhizobium*. QfsR conservation is thus not restricted to Ti plasmids (Figure 1A) but expanded to RepABC plasmids (Table S1).

**Fig.1.**
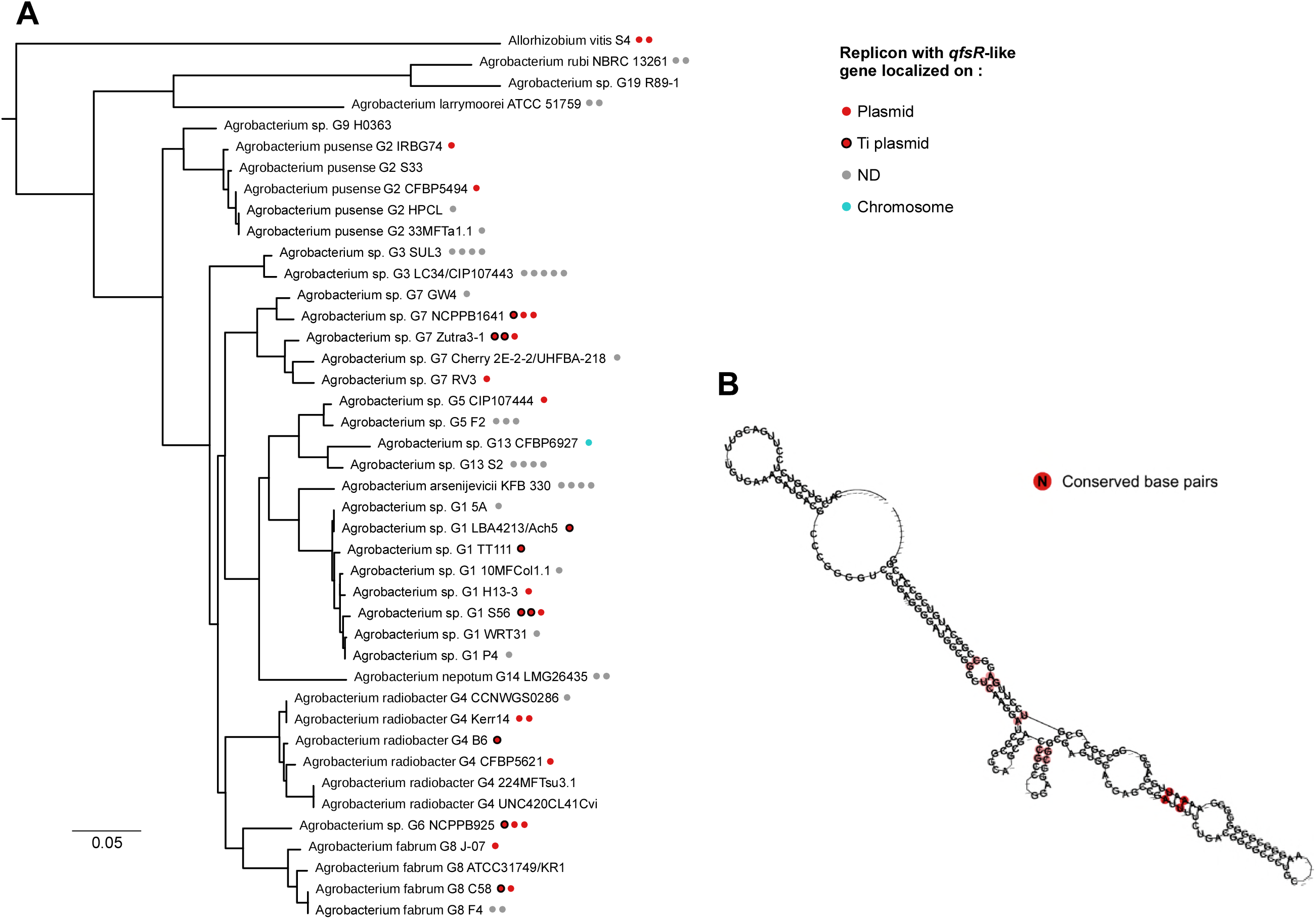
Conservation of QfsR and of its mRNA targets. A) Phylogenetic tree *of Agrobacterium* genomes. The tree was constructed based on an alignment of *recA* sequences, as previously described (62). *qfsR*-like genes are indicated with a disc colored according to replicon-type. B) Consensus structure of the 133 non-redundant QfsR homologs using LocARNA (61).

Results suggested the presence of a *qfsR*-like gene on the other plasmid of the *A. fabrum* C58 strain (pAtC58) (Figure 1A). We also identified four *qfsR* homologs in *Rhizobium etli* bv. *mimosae* MIM1 i.e. one per plasmid (Table S1). Furthermore, two *qfsR* gene homologs could be identified on the same replicon i.e. At plasmid NCPPB925 or plasmid RLG203 from *R. leguminosarum* bv *trifolii* WSM2304. Such a redundancy remains unresolved but is rather widespread since 42 strains harbored at least two putative QfsR homologs and 11 strains presented more than two QfsR homologs (Table S1). We proposed that QfsR belongs to a family of structurally conserved sRNAs, that are transcribed by RepABC-type plasmids.

See also Supplementary Table S1: Occurrence of QfsR and its mRNA targets in bacteria, determined by sequence similarity searches. (Table S1.pdf)

### QfsR is predicted to interact with several polycistronic mRNAs localized on different replicons

Predicting the mRNA targets of QfsR would provide clues for identifying its cellular function. We applied a combination of three algorithms (RNApredator, sTarPicker and IntaRNA) (22)(25)(26)(27) and selected the fifty-four genes identified jointly by them. These candidates were homogeneously distributed among the four replicons, the circular and linear chromosomes and AtC58 and TiC58 plasmids (Table S2). Functional annotation screening using Blast2Go demonstrated an enrichment of the cell motility category. The five highlighted targets were *fliL, flgH, flgI, fliE*, and *flgF*, which belong to a unique polycistronic mRNA encoded from the circular chromosome (Figure 2A, B and C). Five other putative QfsR targets also belong to two polycistronic mRNAs: *exoM* and *exoA* genes are part of the succinoglycan biosynthesis polycistron from the linear chromosome (Figure 2A, D and E) and *trbG, trbL* and *trbK* genes belong to the *traI-trb* Ti plasmid conjugative transfer operon (Figure 2A, F and G). For these ten predicted mRNA targets, the pairing binding site on the QfsR sequence is unique, localized between bases 65 and 107 (Figure 2A, C, E, G). See also Supplementary Table S2: Putative QfsR mRNA targets predicted in common by StarPicker, RNApredator and IntaRNA algorithms. (at the end of the manuscript) *qfsR*-like genes co-occurred with the three polycistrons in most of *Agrobacterium* strains and with *flg-fli* polycistron in *Sinorhizobium* and *Rhizobium* strains (Table S1). To go further, target predictions were performed with QfsR homologs predicted from *Sinorhizobium meliloti* 1021 and *Rhizobium etli* CFN42 genomes. Several putative targets belong to polycistrons among which the *flg-fli* operons like in *A. fabrum* C58 (data not shown).

**Fig.2.**
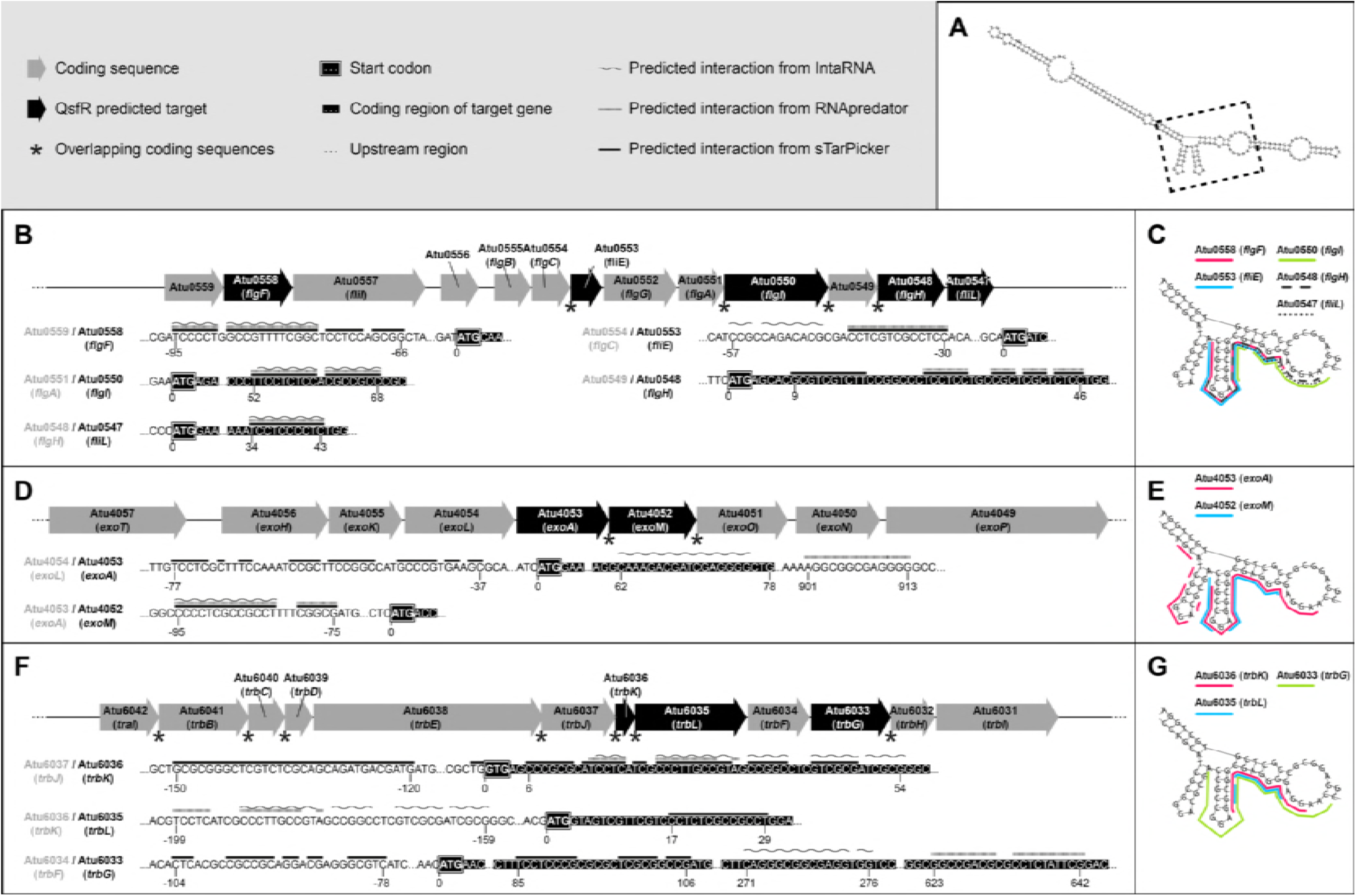
Regions of interaction between predicted target mRNAs and QfsR. A) The predicted secondary structure of QfsR, obtained using the RNAfold algorithm, is shown. B), D), F) Each polycistronic mRNA predicted to interact with QfsR is indicated: black arrows represent coding sequences with a predicted QfsR region of interaction and grey arrows represent coding sequences without a predicted QfsR region of interaction. Overlapping coding sequences are indicated with an asterisk. Predicted interaction regions are highlighted on sequences for each target. B) The data for *flg* polycistronic mRNA. D) The *exo* polycistronic mRNA. F) The *traI-trb* polycistronic mRNA. In C), E) and G), the bases predicted to interact with QfsR putative target mRNAs are indicated on the QfsR secondary structure. It should be noticed that on *flgF, flgI, fliL, exoM, trbK* and *trbL* transcripts, every algorithm predicted identical regions of interaction. Conversely, the regions of interaction on the other transcripts were different according to the algorithms used.

### QfsR interacts directly with mRNA targets by base pairing

To validate the direct interactions between QfsR and its putative mRNA targets, we implemented an *in vivo* reporter system in *E. coli* (28). Each interaction region of the putative mRNA targets (Figure 2) was fused to the superfolder green fluorescent protein gene (sfGFP) in the operon fusion vector pXG30-SF that mimicked a di-cistronic operon. Each construction was co-expressed with QfsR whose gene was cloned under the control of the L-arabinose inducible promoter (pBAD24). The *glmU-glmS* operon fusion in pXG30-SF (29)(28) was used as a control and no variation of fluorescence was detected.

No QfsR-mediated regulation of the fluorescence levels was observed in the *E. coli hfq-deleted* strain (data not shown). Using either wild-type *E. coli* strain or hfq-deleted strain complemented with the *A. fabrum hfq* gene, induction of QfsR production reduced significantly sfGFP fluorescence in strain harboring the *exoA-<u>exoM</u>* reporter fusion, whereas sfGFP fluorescence were significantly induced in strains carrying reporter fusions with *atu0559-<u>flgF</u>, flgC-<u>fliE</u>, flgA-<u>flgI</u> atu0549-<u>flgH</u>, flgH-<u>fliL</u>*, and *trbJ-<u>trbK</u>* (Fig. S2). Furthermore, a change of four nucleotides within the binding site of QfsR (in position 96 to 99: GAGG to AGAA, QfsR*) abolished modulations of fluorescence (Figure 3). Similarly, mutations within the binding site of reporter fusions (exoM*, flgF*, flgI*, fliL* and trbK*) also abolished modulations of fluorescence in the presence of the non-mutated QfsR (Figure 3). However, using compensatory mutations between QfsR* and reporter fusions*, fluorescence levels previously observed were re-established (Figure 3), suggesting the importance of interacting sequence. On the contrary, no variation of fluorescence was detected with the *exoL-<u>exoA</u>* and *trbF-<u>trbG</u>* fusions (Fig. S2). Taken together, these results suggest that QfsR directly interacts within polycistronic mRNAs with the 5’-regions of *exoM, flgF, flgI, fliL* and *trbK* and is likely involved in their post-transcriptional regulation.

**Fig.3.**
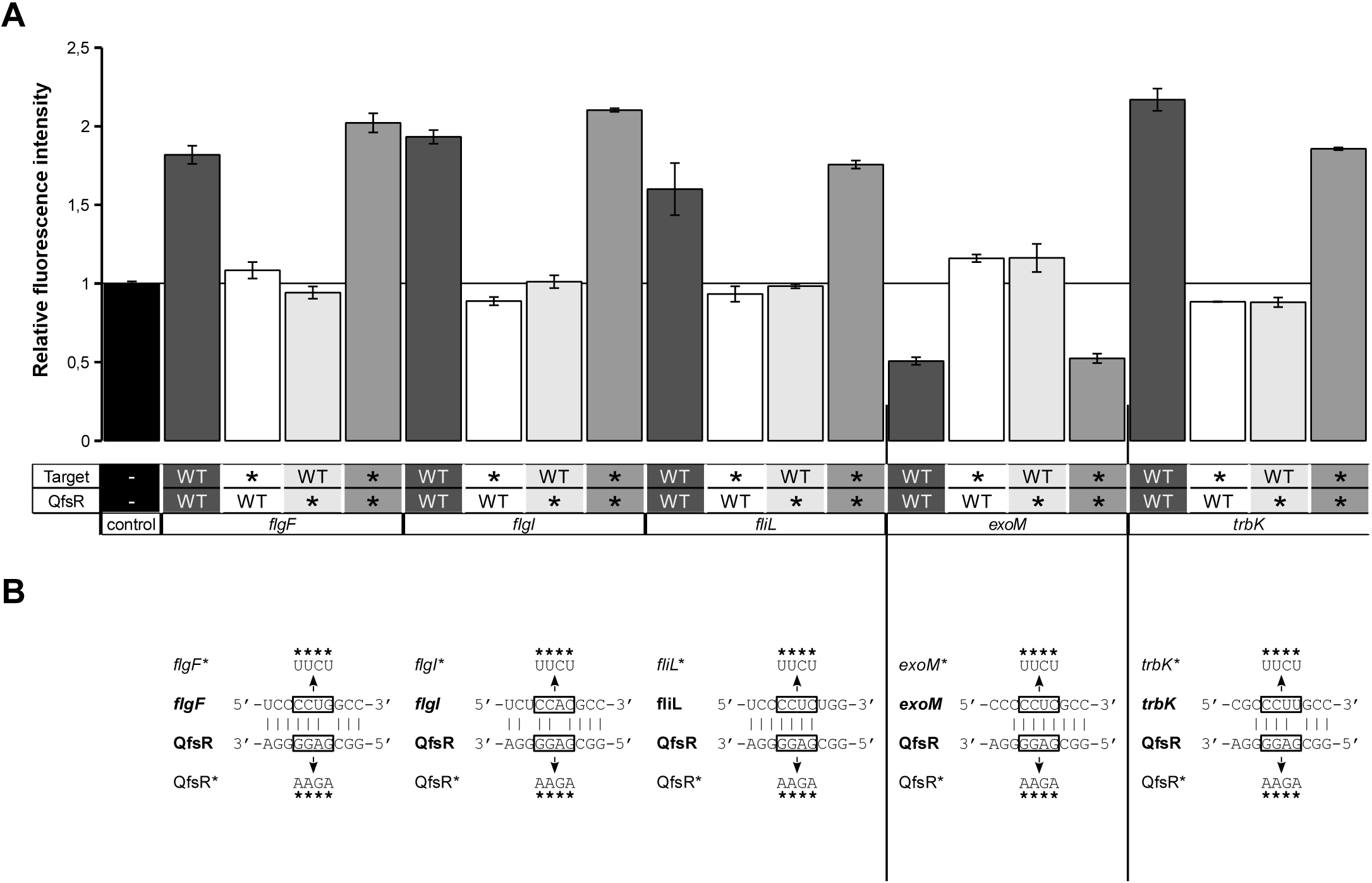
Demonstration of QfsR–mRNA interactions using a heterologous reporter assay. (A) Evaluation of the direct interactions between QfsR and its putative mRNA targets by an *in vivo* reporter system in *E. coli* (28). Relative fluorescence intensity was calculated as the ratio of emitted fluorescence in the presence of QfsR production vs emitted fluorescence in the absence of QfsR (after subtraction of the background fluorescence). Sequences of mRNA targets were fused to the superfolder green fluorescent protein gene (sfGFP) in vector pXG30-SF. Results obtained with *flgF, flgI, fliL, exoM* and *trbK* mRNA targets are presented. The wild-type form and the mutated (QfsR*) form of QfsR were tested as well as mutated forms of mRNA targets (reporter fusions*). Control corresponds to the operon fusion *glmU-glmS* in pXG30-SF as *glmU-glmS* RNA was predicted not to interact with QfsR (29)(28). The 5’ UTR-sgfp fusions for genes *flgH* and *fliE* were tested with the wild-type QfsR only and they are presented in Fig S2. Results presented are those obtained with hfq-deleted strain complemented with the *A. fabrum hfq* gene. Error bars indicate the standard error of the mean. Three independent assays with three technical replicates were performed for each experimental condition. p-value<0.01, by a T-test (B) Predicted interactions of the tested targets with QfsR. Mutations (*) and compensatory mutations are indicated by arrows.

**Fig. S2.**
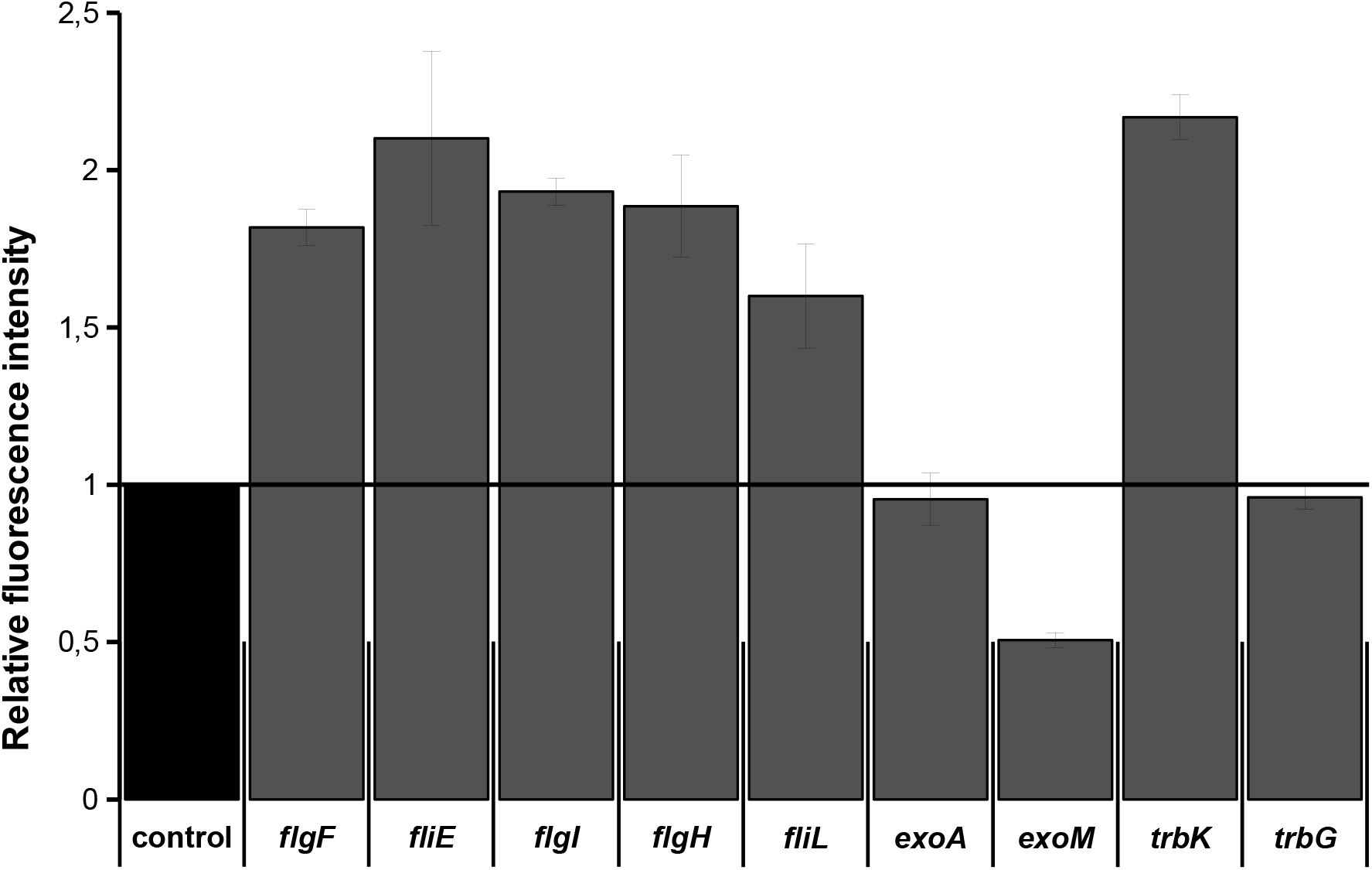
Evaluation of QfsR–mRNA interactions using a heterologous reporter assay. Evaluation of the direct interactions between QfsR and its putative mRNA targets by an *in vivo* reporter system in *E. coli* (28). Relative fluorescence intensity was calculated as the ratio of emitted fluorescence in the presence of QfsR production vs emitted fluorescence in the absence of QfsR (after subtraction of the background fluorescence). Sequences of mRNA targets were fused to the superfolder green fluorescent protein gene (sfGFP) in vector pXG30-SF. The 5’ UTR-sgfp fusions for genes *flgF, flgI, fliL, flgH, fliE, exoA, exoM, trbK* and *trbG* were tested with the wild-type QfsR are presented. Results presented are those obtained with hfq-deleted strain complemented with the *A. fabrum hfq* gene. Control corresponds to the operon fusion *glmU-glmS* in pXG30-SF as *glmU-glmS* RNA was predicted not to interact with QfsR (29)(28). Three independent assays with three technical replicates were performed for each experimental condition. P-value <0.01 by t-test.

### *Deletion of* qfsR *alters motility of* A. fabrum *C58 by modulating mRNA stability of the* flg *polycistron*

To identify the effect of QfsR on *A. fabrum* motility, we generated two strains with either QfsR overexpression (C58/pBBR1MCS-5::QfsR) or knock-out (C58ΔqfsR). We evaluated the swimming ability of these mutants in competition with the wild-type strain (30). Motility indexes (MI) were calculated as the mutant CFU (Colony Forming Unit) number over the wild-type CFU number, normalized by the initial ratio of CFU. As chemotaxis is largely involved in the efficiency of *A. fabrum* motility, assays were performed in the presence or absence of chemo-attractants. Compared to WT, C58ΔqfsR is significantly less motile whatever the presence or absence of any chemo-attractant (Figure 4), suggesting that QfsR activates *A. fabrum* motility independently of chemotaxis.

**Fig.4.**
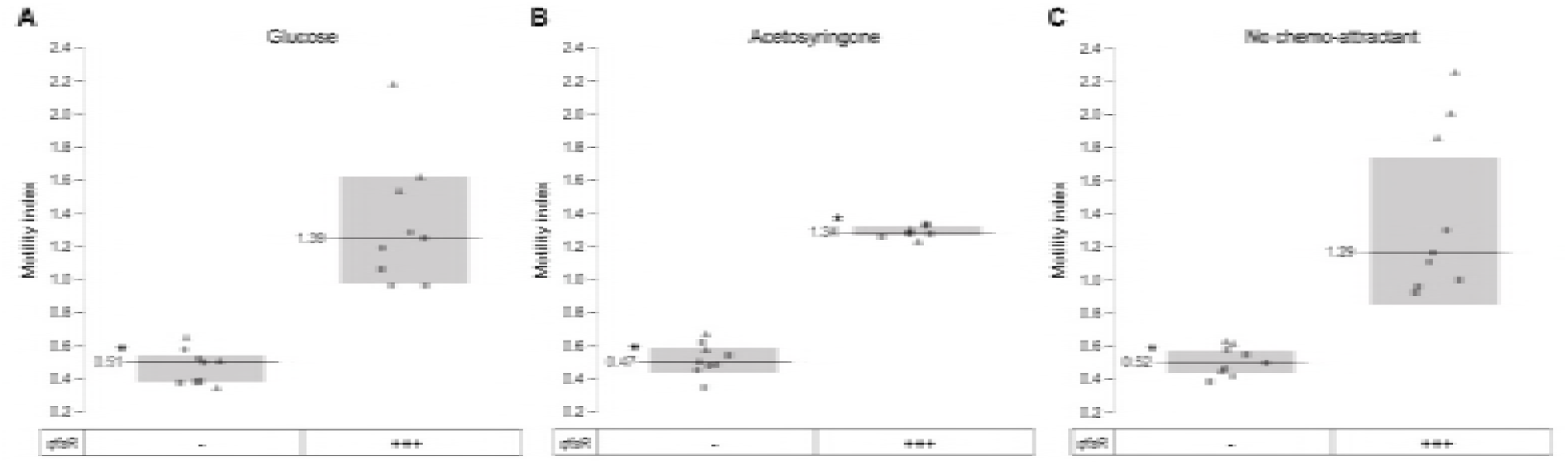
Relative swimming abilities of strains expressing various amount of QfsR. Motility indexes (MI) were calculated as the mutant CFU (Colony Forming Unit) number over the wild-type CFU number, normalized by the initial ratio of CFU. Relative swimming abilities were performed in competition between strains that have different amounts of QfsR. The wildtype was compared either to a strain lacking (-) or to a strain overexpressing QfsR (+++). These competitions were performed in the presence of chemo-attractant glucose (A), or acetosyringone (B), or in the absence of a chemo-attractant, using a sodium-free buffer (C). Statistical significance with a p-value < 0.01 is indicated by an asterisk (T-test). Three independent assays with three technical replicates were performed for each experimental condition.

We then measured expression levels of target genes by qRT-PCR. Whereas QfsR overexpression showed no significant effect on mRNA levels, inactivation of *qfsR* led to a significant decrease of mRNA quantity for all targets of the *flg-fli* operon; mRNA levels were restored to the wild-type levels by ectopic expression of QfsR under its native promoter (Table 1). Similar results were also observed with other genes of this operon not predicted as QfsR targets (*atu0559, flgA* and *flgB*). Furthermore, the mRNA polycistron was less stable in the absence of QfsR as the half-life measured in the wild-type is equal to 4.26 min whereas those of mutant strain is equal to 3.48 min (*fliL* measurement). Taken together, these results suggest that QfsR stabilizes the complete *flg* polycistronic mRNA by interacting directly with the *flg-fli* polycistronic mRNA and the post-transcriptional regulation by QfsR leads to an increase motility of strains harboring Ti plasmid.

**Table 1.** qRT-PCR performed on predicted QfsR mRNA targets in strains harbouring different amounts of QfsR.

| Polycistron | Gene<sup>a</sup> | C58ΔQfsR vs WT |  | C58 / pBBR1MCS-5::QfsR vs WT |  | C58ΔQfsR + QfsR vs WT |  |
| --- | --- | --- | --- | --- | --- | --- | --- |
|  |  | Fold change | p-value<sup>b</sup> | Fold change | p-value<sup>b</sup> | Fold change | p-value<sup>b</sup> |
| *flg* | *atu0559* | 0.53 ± 0.16 | 0.04 | 1.06 ± 0.17 | 0.58 | ND |  |
|  | *flgF<sup>a</sup>* | 0.59 ± 0.01 | 3E-04 | 1.06 ± 0.14 | 0.62 | ND |  |
|  | *flgB* | 0.64 ± 0.12 | 0.04 | 0.95 ± 0.04 | 0.18 | ND |  |
|  | *fliE<sup>a</sup>* | 0.63 ± 0.13 | 0.03 | 1.11 ± 0.16 | 0.42 | ND |  |
|  | *flgA* | 0.52 ± 0.11 | 0.002 | 0.92 ± 0.14 | 0.38 | ND |  |
|  | *flgI<sup>a</sup>* | 0.52 ± 0.03 | 0.002 | 0.93 ± 0.2 | 0.67 | 1.11 ± 0.02 | 0.12 |
|  | *flgH<sup>a</sup>* | 0.55 ± 0.06 | 0.01 | 1.04 ± 0.07 | 0.5 | ND |  |
|  | *fliL<sup>a</sup>* | 0.47 ± 0.03 | 0.002 | 0.86 ± 0.12 | 0.24 | 1.01 ± 0.03 | 0.52 |
| *exo* | *exoT* | 1.36 ± 0.1 | 0.04 | 1.00 ± 0.13 | 0.99 | 0.92 ± 0.04 | 0.24 |
|  | *exoA<sup>a</sup>* | 1.55 ± 0.23 | 0.04 | 0.96 ± 0.15 | 0.76 | 0.96 ± 0.05 | 0.37 |
|  | *exoM<sup>a</sup>* | 1.64 ± 0.19 | 0.04 | 0.98 ± 0.15 | 0.87 | 0.97 ± 0.05 | 0.52 |
|  | *exoN* | 1.48 ± 0.06 | 0.007 | 1.18 ± 0.16 | 0.25 | 0.95 ± 0.05 | 0.26 |

<sup>a</sup> Predicted QfsR mRNA target
<sup>b</sup> p-values were calculated using the student test (3 biological replicates were performed)
ND: Not Determined

**Fig S3.**
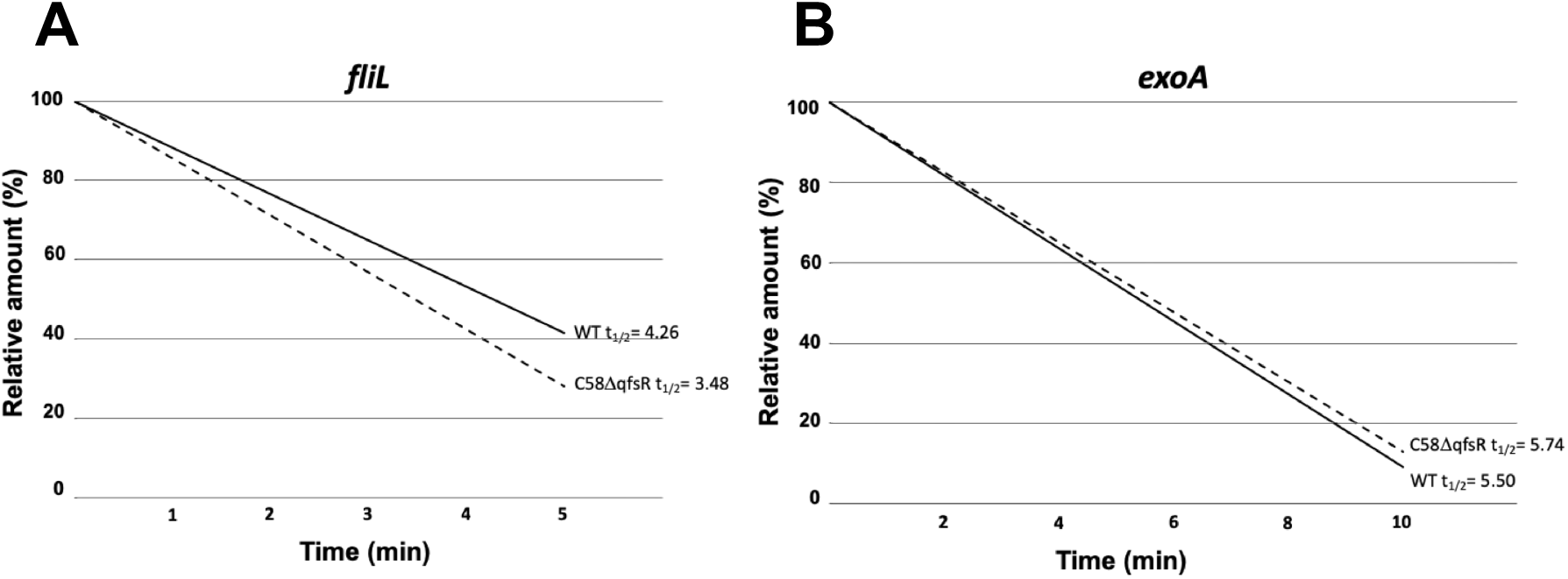
Measurements of transcript half-lives (T_1/2_). A) Transcript stabilities of *flg* operon represented by *fliL* (A) and *exo* operon represented by *exoA* (B) in wild-type and C58ΔqfsR mutant were evaluated by qRT-PCR calculated from at least two biological replicates.

### QfsR represses succinoglycan biosynthesis by modulating mRNA abundance

QfsR is able to directly interact with the 5’ region of *exoM*. Products encoded by the operon to which *exoM* belongs are involved in the biosynthesis of succinoglycan, the most abundant exopolysaccharide produced by *A. fabrum* (31). Although its function remains unclear in *A. fabrum* (32)(33), its glucose linking (β-1,4 and β-1,3) is specifically recognized by Calcofluor staining and can be quantified (34). Whereas WT and overexpressing QfsR strains presented equivalent levels of staining, C58ΔqfsR showed a significant 20% increase in succinoglycan production (data not shown).

To further investigate the underlying mechanism, we evaluated the expression level of *exoM*. A slight but significant increase in *exoM* mRNA abundance was observed in C58ΔqfsR (Table 1). Moreover, expression levels of the other *exo* polycistronic genes not predicted as QfsR targets, i.e. *exoA, exoT, exoN* and *exoP* (Figure 2D), were also increased by the absence of QfsR (Table 1). No difference in mRNA stability of the *exo* polycistron was evidence between WT and C58ΔqfsR (Fig S3). The latter may not be all that surprising considering the slight phenotypic modification.

### Overexpression of QfsR induces Ti plasmid conjugative transfer and quorum sensing signal production

Since QfsR binds to the *trbK* mRNA, we measured the influence of QfsR on the conjugative transfer of pTiC58. Induction of the conjugative transfer requires the presence of agrocinopines. In the absence of these opines, the production of the conjugative apparatus is locked by AccR (Agrocinopine catabolic regulator) that represses the production of the *quorum sensing* transcriptional factor TraR (12). As expected, no detectable transconjugants were observed using WT strain as donor cells in non-inducing conditions, e.g. without agrocinopines (Figure 5A). However, using the strain overexpressing QfsR, transconjugants were obtained (Figure 5A). If the donor cells constitutively expressed the Ti plasmid transfer proteins (due to an *accR* mutation), a high rate frequency of conjugation was measured compared to that obtained with the wild-type strain (Figure 5B). This frequency of conjugation was further enhanced with a donor cell relieved from the AccR repression and overexpressing QfsR (Figure 5B). Furthermore, a change of four nucleotides within the binding site of QfsR (QfsR*) suppressed the increase in the number of transconjugants observed when QfsR is overexpressed (Fig. S4). Overall, QfsR, when present in excess, plays an activator role on the conjugation of the Ti plasmid. This QfsR regulation involves its direct binding site and likely its direct interaction with *trbK*.

**Fig.5.**
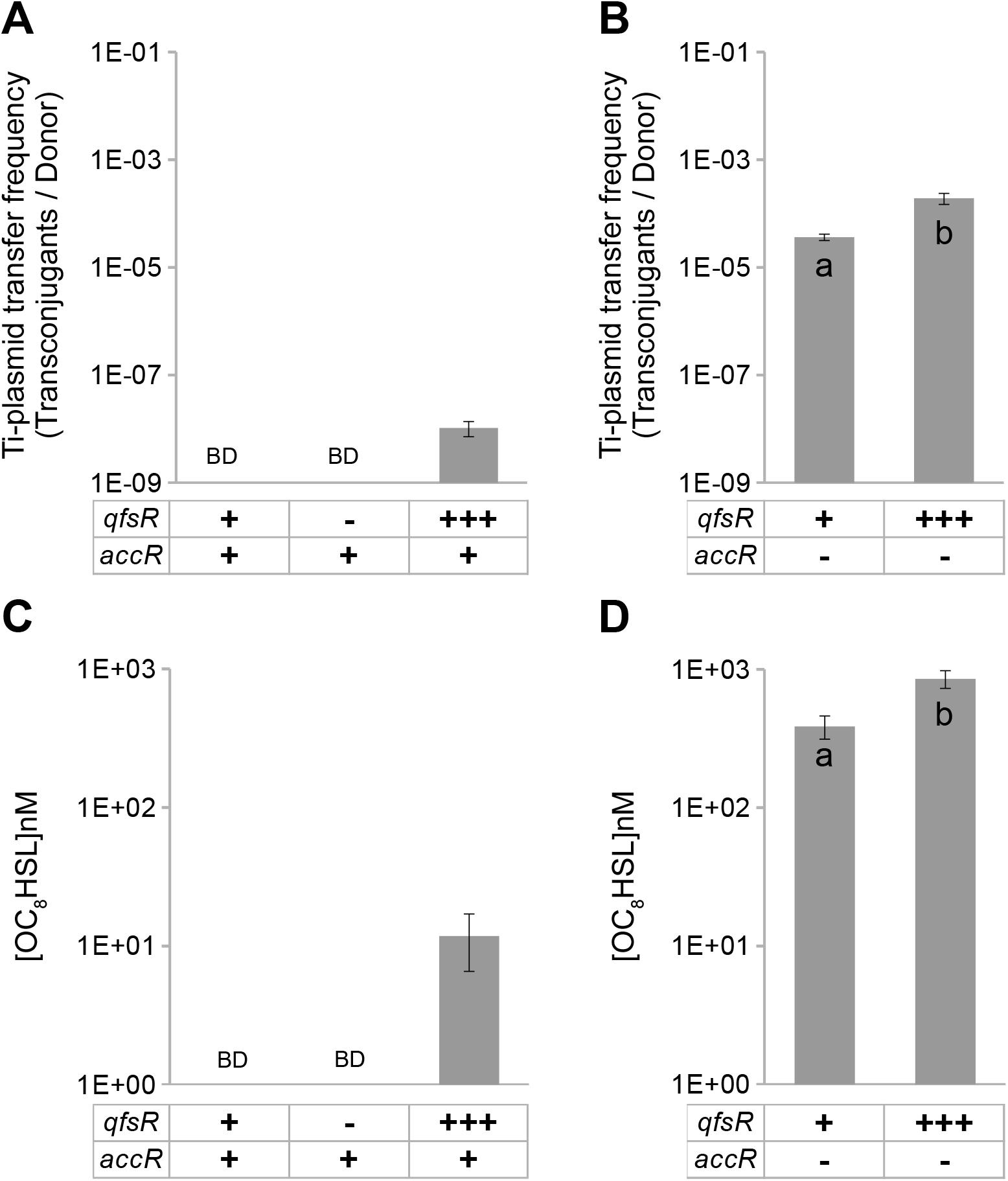
Quantification of Ti plasmid conjugative transfer and quorum sensing signals. A) Transfer frequencies *of A. fabrum* C58 pTi::Km (AT296), pTi lacking QfsR (AT193), or pTi::Km in a strain overproducing QfsR were evaluated *in vitro* (AT298). B) Transfer frequencies of *A. fabrum* C58 pTiaccR::Gm, with or without the overproduction of QfsR (AT315 and AT314), were evaluated *in vitro* (logarithm scales). C) *In vitro* quantification of the OC_8_HSLs produced by the *A. fabrum* C58 wild type strain harbouring different amounts of QfsR or D) by a strain lacking AccR, with or without the overproduction of QfsR (logarithm scales). BD corresponds to ‘below detection’, i.e < 10^−09^ for transfer frequency and < 1 pmol.g^−1^ for OC_8_HSL production. Three independent assays were performed for conjugative transfer and quorum sensing signal quantifications. P-value ≤ 0.01by a T-test.

**Fig. S4.**
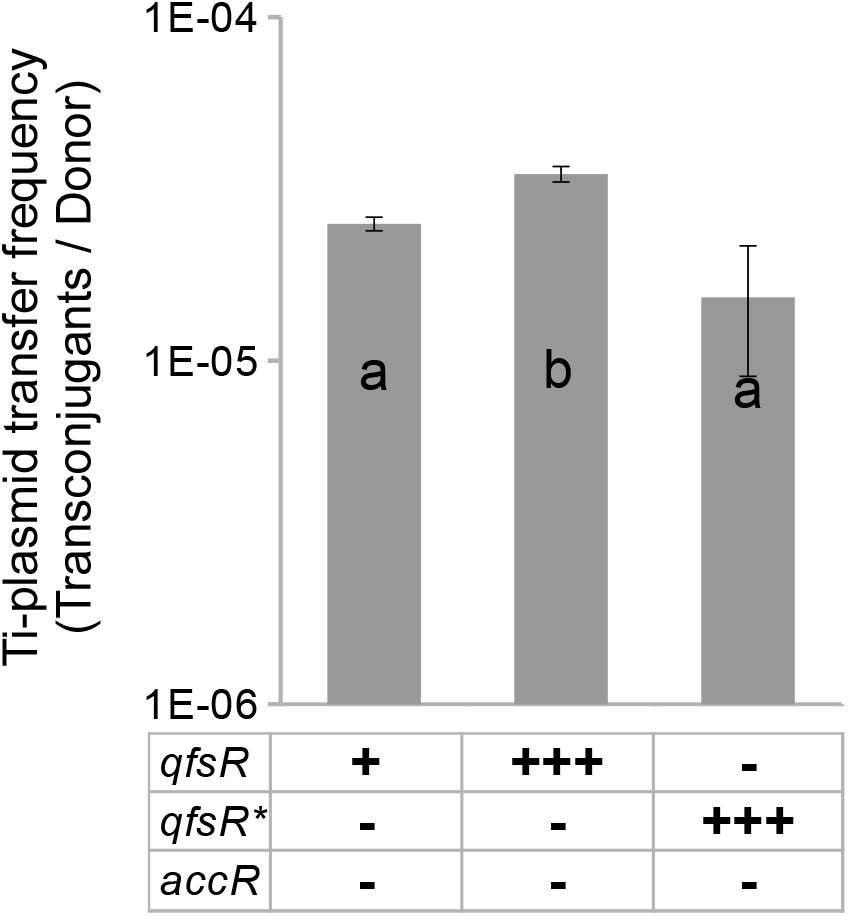
Quantification of Ti plasmid conjugative transfer. In vitro evaluation of transfer frequencies of *A. fabrum* C58 pTiaccR:: Gm, without or with the overproduction of QfsR WT or with the overproduction of QfsR harbouring a change of four nucleotides within its binding site (QfsR*) (logarithm scales). Three independent biological assays with three technical replicates were performed for each experimental condition. P-value ≤0.01 by t-test.

The first gene of the *traI-trb* polycistron (containing *trbK)* encodes TraI, which produces the *quorum sensing* signal, OC_8_HSL. OC_8_HSL is required to stabilize TraR and thus to allow the production of the conjugative apparatus (12). Quorum sensing signal was not detected in the WT whereas OC_8_HSL was detected when QfsR is overexpressed (Figure 5C). Furthermore, accumulation of OC_8_HSL was significantly higher in an *accR* mutant background overexpressing QfsR compared to that measured in the *accR* mutant (Figure 5D). Overall, these results showed that OC_8_HSL production increases when QfsR is overexpressed. As *traI* belongs to the *traI-trb* polycistron (12), we hypothesized that QfsR stabilizes *traI* mRNA, and maybe the complete polycistronic mRNA by its direct interaction with *trbK*.

### QfsR levels increase under acidic environment

As QfsR amount is important for its regulatory functions, we measured QfsR expression levels under different growth conditions by qRT-PCR. Except for acidic environment (AB medium at pH 5.6) where QfsR expression increased by a level of 8-fold (data not shown), QfsR seems to be constantly expressed in the other growth conditions tested (exponential vs stationary phases, minimal vs rich medium, tumor and plant extracts). It should also be noticed that Ti plasmid copy number raises under virulence condition (15)(17)(14)(16) and in accordance expression levels of QfsR, whose gene is localized on the Ti plasmid, increased at levels equivalent to the increase of pTi copy number (around 4 fold).

## Discussion

### sRNAs regulate motility, biofilm formation and quorum sensing

To date, a surprisingly large number of sRNAs modulate directly or indirectly quorum sensing (39), motility and/or biofilm formation (40). In several gamma-proteobacteria, the Csr/Rsm system involves sRNAs that regulate virulence, biofilm, motility and quorum sensing together by titrating the CsrA/RsmA protein, which acts to regulate genes at a post-transcriptional level (38). In *E. coli* and *Salmonella*, several *trans-acting* sRNAs regulate motility and/or biofilm formations by their direct interaction with mRNAs of master regulators i.e. FlhDC, CsgD and RpoS (41). In the present study, we demonstrated QfsR to be a *trans-acting* sRNA that directly regulates the accumulation of mRNA-encoding structural proteins of flagella, enzymes involved in exopolysaccharides biosynthesis and quorum sensing signal synthesis (Figures 3, S2, Table 1). Altogether, QfsR seems to function as a hub connecting regulatory circuits of flagella, exopolysaccharides and quorum sensing signal (42).

Crosstalk may occur between QfsR mRNA targets. RNA-RNA interactions between QfsR and its mRNA targets use an identical base pairing site in QfsR (Figure 2C, E, G). This finding led to hypothesize a competition for QfsR and cross-regulation between QfsR mRNA targets (43)(35). Although the affinities between QfsR and its targets and the fate of QfsR after base pairing are not known, it has been established that an artificial increase in QfsR has no effect on the *flg* transcript but increases pTiC58 conjugative transfer and OC_8_HSL biosynthesis (Table 1, Figure 5). These data suggest a saturation of the QfsR binding sites on *flg* transcripts but not on *traI-trb* transcripts. Based on these data, we hypothesized a crosstalk between conjugation and motility that might be beneficial for Ti plasmid maintenance and dissemination. Indeed, conjugative transfer occurs in the presence of recipient bacteria and in their absence, donors must shift to new environments where recipient cells are present. The control exerted by QfsR on the regulation of motility may allow pTi dissemination by conjugation.

### QfsR exerts a feedforward control on pTiC58 conjugation

The expression of pTi transfer genes is strictly controlled by transcriptional regulators. TraR is absolutely required to activate the production of the conjugative apparatus and its expression is locked by the repressor AccR. This repression is relieved by agrocinopine (12). TraR forms a complex with OC_8_HSL and TraR without its ligand OC_8_HSL is insufficiently stable to be efficient, especially at the initiation of conjugation or in the absence of agrocinopines (44)(45)(46). To be triggered, the mechanism requires at least a small amount of available OC_8_HSL for the formation of the initial TraR-OC_8_HSL complex. OC_8_HSL is produced by TraI, encoded by the first gene of the *traI-trb* polycistron (12), whose expression is induced by the TraR-OC_8_HSL complex. Until now, the mechanism by which OC_8_HSL production is initiated remains to be elucidated. Production of OC_8_HSL and pTi conjugative transfer were detectable when QfsR was overexpressed (Figure 5), suggesting a stabilization of the *traI-trb* operonic mRNA by QfsR. Thus, an increase in QfsR amount could allow the stabilization of *traI-trb* mRNA, leading to the initial production of OC_8_HSL. As QfsR increased in acid environment and condition triggering virulence i.e. before the production of TraR, we proposed QfsR as a feedforward modulator of Ti plasmid conjugative transfer.

QfsR exerts the posttranscriptional regulation of polycistronic mRNAs. QfsR represses *exo* polycistronic mRNA and activates *flg* and *traI-trb* polycistronic mRNAs (Table 1, Figure 3). Whatever the targets, QfsR directly interacts within the polycistronic molecule and coordinates expression of its encoded genes (Table 1, Figure 2), highlighting the involvement of QfsR in the regulation of polycistronic mRNAs (49). Additional studies are necessary to decipher the mechanism involved. The advantage of such RNA-RNA interactions should be analyzed according to the characteristics shared between these polycistrons. In particular the presence of coding sequences with short distances between the stop codon of the upstream gene and the start codon of the downstream gene and even overlapping coding sequences should been taken into consideration (Figure 2). This implies that mRNA molecules should remain uncleaved for correct translation. QfsR may be involved in the maintenance of polycitronic mRNA integrity until translation.

### Plasmid-transcribed QfsR regulates chromosomal targets

So far, only indirect evidences have proposed that some horizontally acquired sRNAs regulate directly core genome elements (50). Here, we present strong evidences that QfsR regulates directly chromosome-encoded mRNAs, which highlights crosstalk between replicons. Moreover, there are only a few plasmid regulators controlling chromosomal genes, all of them being DNA binding proteins (6)(7)(8). Ti plasmids are conjugative elements, meaning that they are present in various *Agrobacterium* genomic backgrounds that could differ in sequence binding sites with QfsR (51). This finding raised questions concerning the conservation of QfsR regulatory functions since QfsR-C58 regulation could be efficient in *Agrobacterium* strains only if target genes harbor conserved binding sites. Interestingly, it has been observed that significant conservation for RNA-RNA binding sites and corresponding short sequence stretches are highly and specifically conserved in four of the five QfsR chromosome-encoded targets among *Agrobacterium* strains (data not shown)(52)(53).

Analyses of sequence and structure conservations led to the hypothesis that QfsR is widely conserved among RepABC plasmids (Figure 1, Table S1). Although the evolutionary forces that shape sRNAs are not yet understood, sRNAs are described to evolve rapidly (53). sRNA-target pairs conserved between *E. coli* and *S. enterica* are not found in general across more distantly related species (52)(53). Consistently, we failed to identify orthologous QfsR mRNA targets in other *Rhizobiaceae*. However, the *flg* polycistrons were predicted as QfsR homolog targets in *Sinorhizobium* and *Rhizobium* but target genes inside the *flg* polycistrons and the localization of binding sites are different. This illustrates (i) plasmidic sRNAs as driving force for regulatory turnover and even regulatory novelty and (ii) ecological advantage for the regulation of motility by the QfsR family.

In conclusion, QfsR is, to our knowledge, the first plasmid-encoded sRNA described to regulate chromosomal polycistronic mRNAs via a direct base pairing interaction. Such replicon crosstalk using plasmidic sRNA and chromosome mRNAs could be widespread events.

## Materials and Methods

### Bacterial strains, media and growth conditions

Bacterial strains used in this study are listed in Table S3. *Escherichia coli* strains were grown, with shaking (160 rpm), at 37°C in Luria-Bertani media or in M9 media supplemented with 0.4% glycerol and 0.2% casamino-acid (54). *A. fabrum* strains were grown, with shaking (160rpm), at 28°C in YPG-rich media (yeast extract 5 g.L^-1^; peptone 5 g.L^-1^; glucose 10 g.L^-1^; pH=7.2) or modified AB induction media (1 g N⅛Cl, 0.3 g MgSO_4_·7H_2_O, 0.15 g KCl, 0.01 g CaCl_2_, 2.5 mg FeSO_4_·7H_2_O, 2 mM phosphate buffer (pH 5.6), 50 mM 2-(4-morpholino)-ethane sulfonic acid (MES), 0.5% glucose per L, pH 5.6), with or without 200 μM.mL^-1^ acetosyringone (200 μM) (55). Media were supplemented, when necessary, with antibiotics at the following concentrations: 100 μg.mL^-1^ ampicillin, 5 μg.mL^-1^ gentamicin, 20 μg.mL^-1^ chloramphenicol, 10 μg.mL”^1^ tetracycline and 50 μg.mL^-1^ kanamycin for *E. coli*, and 25 μg.mL^-1^ gentamicin, 25 μg.mL^-1^ neomycin, 25 μg.mL^-1^ kanamycin and 100 μg.mL^-1^ spectinomicin for *A. fabrum*.

### Bacterial strain constructions

The DNA fragment between nucleotides 139,052 and 139,269 of the Ti plasmid (corresponding to *qfsR* gene) was amplified, by PCR, using primers 182 and 183 (Table S3). Fragments were purified and digested using *SalI* and *BamHI* enzymes and cloned into pBBR1MCS-5 and pBBR1MCS-2, giving rise to pARA007 and pARA006, respectively (Table S3). These plasmids were electroporated into the *A. fabrum* C58 strains, giving rise to AT204 (C58/pBBRMCS-5::QfsR) and AT315 (C58/pTiaccR::Gm/pBBRMCS-2::QfsR) respectively. An increase of QfsR level of about 10 fold was measured in these strains. *A. fabrum* C58 strain was used to construct a deleted mutant, as previously described (56). The genomic region between bases 138,991 and 139,368, including *qfsR*, was deleted and replaced by the neomycine-kanamycine resistance gene *nptII*, giving the strain AT193 where the QfsR level was equivalent to the background level (Table S3). The introduction of *qfsR* gene under the control of its own promoter (ectopic expression) in AT193 allowed an expression of QfsR similar to the level observed in the wild-type strain. Mutant strains showed no distinctive phenotypes, they grew similarly to the wild type and showed no morphological specificity.

### Determination of RNA 5 ‘- and 3’-ends by RACE-PCR

The 5’ and 3’-end of QfsR were determined by RACE-PCR, as previously described (22). Reverse transcription was performed with the gene specific primer A311 (Table S3) and the junctions between the 5’-end and the 3’-end of the RNA were amplified using primers 295 and 296 (Table S3). The resulting RACE-PCR products were cloned into the pGEMT-easy. DNA fragments were then sequenced using the M13fwd and the M13rev primers (Invitrogen).

### Quantitative RT-PCR analysis (qRT-PCR)

Primers used are listed in Table S3. For each strain, independent triplicate *A. fabrum* cultures were grown in LPG media. RNA extraction, reverse transcription and quantitative PCR were performed, as previously described (22), using the SuperScript III (Life Technologies) and the DyNAmo Flash SYBR Green qPCR kit (Thermo scientific) with the LC480 Lightcycler from Roche. The specificity of the PCR primers was verified with a melting curve analysis.

### Measurement of mRNA decay

An overnight culture was diluted to OD_600_=0.05 and cells were grown to OD_600_=0.4. A reference sample was drawn from culture and then rifampicin was added to arrest transcription (final concentration = 250 μg/mL). After rifampicin treatment, cells were harvested from culture at 5 and 10 min. To preserve cellular RNA intact, culture samples were mixed with one-tenth volume of a “stop-solution” composed 10% buffer-saturated phenol in ethanol and then chilled rapidly. RNA were extracted as described previously (22), reverse transcribed and expression level of gene of interest *(flgF, flgI, flgH, fliL, exoA, exoM* and *exoT*) were evaluated by quantitative PCR.

### eGFP-mediated fluorescence constructs and assays

To construct an L-arabinose inducible QfsR, the *qfsR* gene was PCR amplified (between bases 138,883 and 139,451 of the Ti plasmid) using primers 332 and 333 (Table S3), with a sense primer designed such that it started with the QfsR +1 site and a HindIII site was added to its sequence. The antisense primer (333) binds ~ 189 nt downstream of the *rna1083* terminator and adds an XbaI site to its sequence. The PCR products were ligated into pBAD-24 (Invitrogen) previously digested with XbaI and HindIII, yielding plasmid pBAD24::QfsR. The plasmid pXF30-SF served as the backbone for the construction of plasmids with translational sfGFP fusions (28). Each fragment of interest was amplified by PCR (length around 300bp) and cloned between NheI/NsiI restriction sites in order to induce, with anhydrotetracycline, the expression of the 5’-UTR translation fusions of the predicted target genes. Cloning fragments correspond to sequences of the circular chromosome between bases 544,628 and 544,895 for *flgF*, between bases 540,955 and 541,184 for *fliE*, between bases 539,271 and 539,488 for *flgI*, between bases 537,620 and 537,855 for *flgH*, and between 536,877 and 537,136 for *fliL*. Sequences of the linear chromosome were between bases 1171,004 and 1171,263 for *exoA* (which corresponds to region of interaction predicted by sTarPicker and IntaRNA) and between bases 1169,976 and 1170,274 for *exoM*, and cloning fragments correspond to sequences of the Ti plasmid between bases 46,228 and 46,463 for *trbK* (which corresponds to the complete region of interaction predicted by RNApredator and IntaRNA but only a part of the region of interaction predicted by sTarPicker) and between bases 44,083 and 44,348 for *trbG* (which corresponds to the region of interaction predicted by sTarPicker). *E. coli* derivates (TOP10F’, TOP10F’ *hfq*::kan, TOP10F’ *hfq*::kan/pBBGsyn_hfq) containing pBAD24::QfsR and plasmid pXF30-SF, with translational fusions between target mRNA and sfGFP, were grown overnight in M9 medium supplemented with 0.4% glycerol and 0.2% casamino acids. Three independent biological replicates were performed for each strain. Twice-washed bacteria were then resuspended into fresh M9 medium supplemented with 0.4% glycerol and appropriate amino-acids. After bacterial growth, 3 technical replicates of 200 μL aliquots, at OD_600_ = 0.8, were transferred to a 96-well microtiter plate and incubated at 37°C, with shaking, in the microplate reader Infinite^®^ 200 PRO (Tecan). When necessary 0.2% arabinose was added to induce QfsR expression. After 2 hours incubation, anhydrotetracycline (0.5μg.mL^-1^) was added for the induction of translation fusion expression. Fluorescence was read on every 10 min during a total of at least 6 hours. Readings were taken ten times during a 20 ms period with wavelength at 485 nm for excitation and 530 nm for emission. Values were normalized to the culture OD_600_ and measurement of fluorescence prior to anhydrotetracycline induction evaluated the autofluorescence levels, which were then substracted. The effect of QfsR on the translation fusions was measured as the ratio of the values obtained with and without L-arabinose in the culture media.

### *Conjugation assays* in vitro

Conjugation assays were performed, as previously described, in an equal ratio (1:1) (37). Suspension dilutions of conjugation assays were spotted onto selective agar media. Different bacterial populations were, thus, enumerated. Transfer frequencies were evaluated as the number of transconjugants over the number of donors. Three independent biological assays with three technical replicates were performed for each experimental condition.

### Quantification of OC_8_HSL

OC_8_HSL was quantified, as previously described, using the OC_8_-HSL-bioindicator strain *A. tumefaciens* NTLR4 (57). OC_8_HSL concentrations were evaluated by comparison with a calibration curve obtained from pure OC_8_HSL, using the ImageJ software (58). Three biological replicates were performed.

### Calcofluor fluorescence quantification of culture supernatants

Calcofluor fluorescence quantifications were performed, as previously described (59), with the following modifications: bacteria were grown in AB medium supplemented with 0.4% mannitol and 3.5 μL of White stain calcofluor, at 1 mg.mL^-1^ (Sigma-Aldrich), were added to 196.5 μL of frozen culture supernatant in a 96-well plate. Fluorescence was read on the microplate reader Infinite^®^ 200 PRO, from Tecan, with excitation at 365 nm and emission at 435 nm. Readings were taken ten times during a 20 ms period. The appropriate blank was subtracted from each row of fluorescence readings and values were normalized to the OD, at 600 nm, of the bacterial cultures. Experiments were performed in triplicates.

### Chemotaxis assays

Chemotaxis assays were performed, as previously described, with either acetosyringone or glucose (YPG was used as glucose containing medium) as chemo-attractants or without any chemo-attractant in sodium-free buffer (10 mM KPO4 and 1 mM EDTA) (30). Briefly, 10 mL of exponentially grown bacteria were washed twice in sodium-free buffer and then concentrated in 3 mL. Capillary assays are performed in competitions in an equal ratio 1:1. Suspension dilutions of chemotaxis assays were spotted onto selective agar media. Different bacterial populations were, thus, enumerated. Motility indexes were calculated as the ratio of the number of mutant strains versus the number of wild-type strains. Three independent biological assays with three technical replicates were performed for each experimental condition.

### Statistical analyses

Normal distributions were tested with Shapiro-Wilk test. For comparisons between the groups for the determination of (i) *gfp* reporter fusions, (ii) motility indexes (iii) and conjugative transfer, independent T-test was employed (p-value<0.05). For the statistical analysis of qRT-PCR a Student test was used (p-value<0.05).

### Secondary structure predictions and similarity searches

RNA secondary structures were predicted using the RNAfold algorithm. Webserver default options were chosen for folding. Sequence similarity searches were performed using the BlastN algorithm with BLAST+ 2.5.0 software (60). Sequence similarities were searched in the NCBI Nucleotide collection (nt) and MicroScope genome database (23). Multiple alignment of non-redundant RNA molecules were performed to obtain a consensus secondary structure using LocARNA (61). RNAforester was then used with the local similarity model to select QfsR homolog according to their secondary structure conservation (24).

### Bacterial small RNA target predictions

Target genes of QfsR were predicted from the *A. fabrum* C58 genome using IntaRNA (27), sTarPicker (26) and RNApredator (25) algorithms. Targets of interest were selected as previously described (22).

### Gene Ontology enrichment analysis

*A. fabrum* Blast2Go annotation, with default settings, was used to perform an enrichment analysis with Fisher’s exact test on QfsR putative target genes.

## Acknowledgements

This study was supported by the PEPS INSB and PEPII INSB-INSMI-INS2I “Bio-Math-Info” funding from the Centre National de la Recherche Scientifique. M. Dequivre and B. Diel received doctoral grants from the French Ministère de l’Education nationale de l’Enseignement Supérieur et de la Recherche. The authors thank Dominique Loqué, William Nasser and Sylvie Reverchon for comments on the manuscript, Xavier Nesme, Denis Faure, Cristina Vieira and Yvan Rahbé for helpful discussion, Eliane Hajnsdorf for providing us the *E. coli* Hfq mutant strain, and Jörg Vogel for providing us with pXF30 plasmid. The authors are grateful to Camille Villard and Agnès Rodrigue for helpful technical assistance. qPCR experiments were performed at the DTAMB platform from the BioEnviS scientific federation.

---

**Table S2.** Putative QfsR mRNA targets predicted in common by StarPicker, RNApredator and IntaRNA algorithms.

| Accession number | Gene name | Encoded Function | Rank of prediction | Identical region of interaction for the three algorithms onto QfsR | Identical region of interaction for the three algorithms onto the mRNA target |
| --- | --- | --- | --- | --- | --- |
| **Circular chromosome** |  |  |  |  |  |
| Atu0048 |  | Unknown function | 23 | X | X |
| Atu0462 |  | Peptidase m15 | 32 | X | X |
| Atu0510 | *phaD* | Monovalent cation H<sup>+</sup> antiporter subunit b | 22 |  |  |
| Atu0547 | *fliL* | Flagellar basal body protein | 10 | X | X |
| Atu0548 | *flgH* | Flagellar L-ring protein | 11 | X |  |
| Atu0550 | *flgI* | Flagellar P-ring protein | 3 | X | X |
| Atu0553 | *fliE* | Flagellar hook-basal body protein | 53 | X | X |
| Atu0558 | *flgF* | Flagellar basal body rod protein | 21 | X | X |
| Atu0602 | *glnA* | Glutamine synthetase | 9 | X | X |
| Atu0648 |  | Transcription accessory protein | 4 | X | X |
| Atu0669 |  | Unknown function | 8 | X | X |
| Atu0712 |  | ATP-binding protein | 41 | X |  |
| Atu0766 | *ialB* | Invasion protein | 37 | X | X |
| Atu0847 | *ohr* | Organic hydroperoxide resistance protein | 50 |  |  |
| Atu0936 |  | Methyltransferase | 45 |  |  |
| Atu0956 | *gp35* | Primosomal replication protein N | 38 |  |  |
| Atu0990 |  | 3-oxoacyl-ACP reductase | 15 | X | X |
| Atu1057 | *nudF* | ADP-ribose pyrophosphatase | 13 | X | X |
| Atu1222 |  | Hypothetical protein | 51 |  |  |
| Atu1764 | *metF* | Methylenetetrahydropholate reductase | 6 | X | X |
| Atu1791 |  | Sugar ABC transporter permease | 18 | X | X |
| Atu1916 |  | Periplasmic protein | 12 | X | X |
| Atu1978 |  | Hypothetical protein | 20 | X | X |
| Atu1986 |  | Histidine kinase | 14 | X |  |
| Atu2114 | *djlA* | Molecular chaperon | 49 | X |  |
| Atu2248 |  | Hypothetical protein | 39 | X |  |
| Atu2316 |  | Membrane protein | 30 | X |  |
| Atu2380 |  | Globin | 40 | X | X |
| Atu2389 |  | ABC transporter permease | 1 | X | X |
| Atu2420 |  | ATPase | 42 | X |  |
| Atu2515 | *dppD* | Peptide ABC transporter ATP-binding protein | 5 | X | X |
| Atu8181 |  | Hypothetical protein | 19 | X |  |
| Atu2655 |  | E3 ubiquitin-protein ligase | 36 | X | X |
| **Linear chromosome** |  |  |  |  |  |
| Atu3166 | *scrK* | Sugar kinase | 24 |  |  |
| Atu3214 |  | Peptide ABC transporter permease | 27 | X |  |
| Atu3530 | *uxuB* | Mannitol dehydrogenase | 31 | X | X |
| Atu3548 |  | Glycosyl transferase family 1 | 47 | X | X |
| Atu3886 | *pldB* | Lysophospholipase | 7 | X | X |
| Atu3920 |  | Hypothetical protein | 48 | X |  |
| Atu3975 | *glf* | UDP-galactopyranose mutase | 16 | X |  |
| Atu4052 | *exoM* | Glycosyl transferase family A | 25 | X | X |
| Atu4053 | *exoA* | Succinoglycan biosynthesis protein | 26 | X |  |
| Atu4114 | *dppB* | Peptide ABC transporter permease | 29 | X | X |
| Atu4222 | *soxA* | Sarcosine oxidase alpha subunit | 2 | X |  |
| Atu4245 |  | ABC transporter permease | 44 | X | X |
| Atu4642 | *katG* | Peroxydase | 43 | X |  |
| Atu4772 |  | DNA-binding response regulator | 17 | X | X |
| **At plasmid** |  |  |  |  |  |
| Atu5084 |  | Transcriptional regulator | 46 | X | X |
| Atu5164 | *avhB3* | Type IV secretion protein 3 | 28 | X |  |
| Atu5514 | *dnaJ* | Molecular protein | 33 | X | X |
| **Ti plasmid** |  |  |  |  |  |
| Atu6033 | *trbG* | Conjugal transfer protein | 54 | X |  |
| Atu6035 | *trbL* | Conjugal transfer protein | 35 | X |  |
| Atu6036 | *trbK* | Entry exclusion protein (conjugal transfer protein) | 34 | X | X |
| Atu6053 |  | Hypothetical protein | 52 | X |  |

**Table S3.**
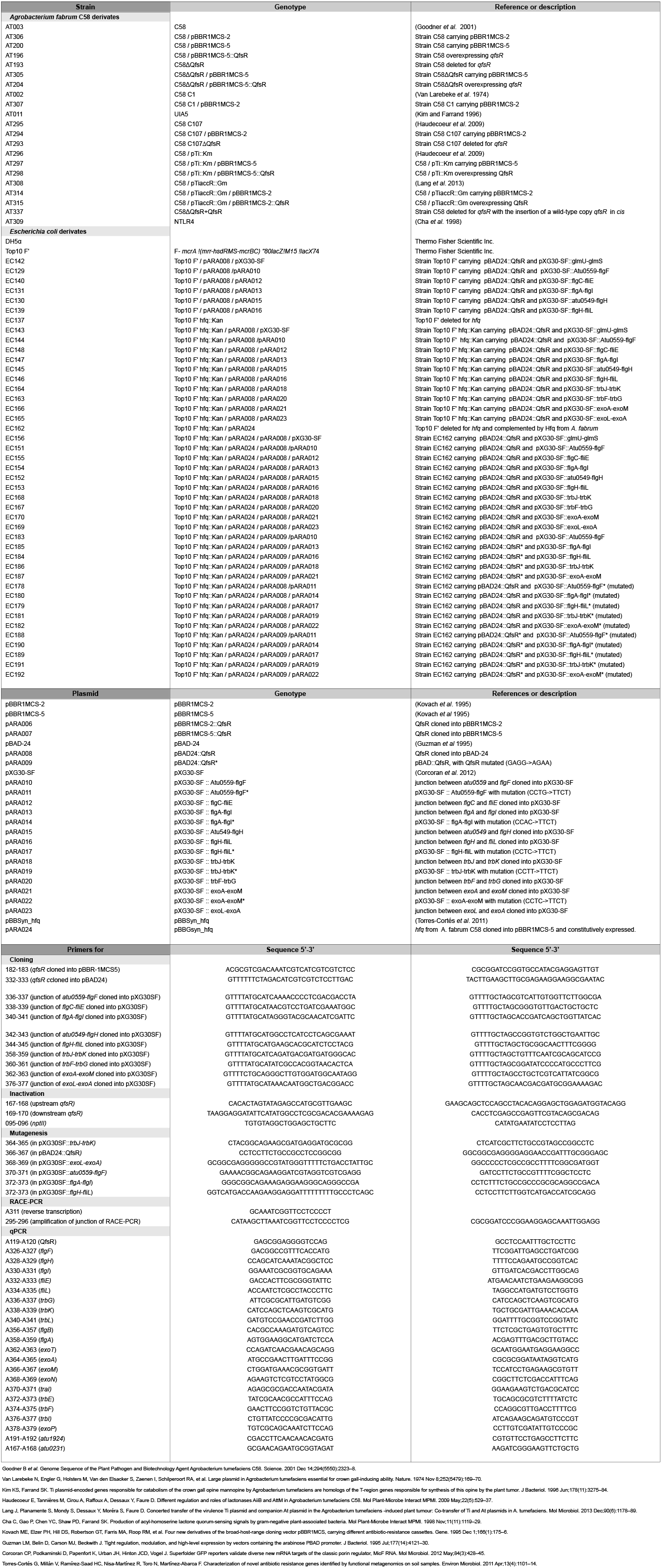
Bacterial strains, plasmids and oligonucleotides used in this study.

